# Apolipoprotein-based regulation of ganglioside metabolism upon secretase activity inhibition in iPSC-derived cerebral organoids

**DOI:** 10.1101/2023.08.30.555505

**Authors:** Durga Jha, Tereza Váňová, Dáša Bohačiaková, Zdeněk Spáčil

**Author notes:** Corresponding Author: Dr. Zdenek Spacil, Masaryk University, Faculty of Science, RECETOX, Kamenice 753, 625 00 Brno, Czechia, E‑mail or.

## Abstract

Beta and gamma-secretase inhibitors have been of pharmacological interest to reduce abeta (Aβ) formation and aggregation, one of the defining characteristics of Alzheimer’s disease (AD). Recent research indicates that Apolipoprotein E (ApoE), a genetic risk factor for AD, can regulate secretase activity. Secretase inhibitor-induced elevation of neuronal membrane lipids has been documented in 2D models. Due to their enhanced ability to reproduce AD-like pathology and ease of performing the experimental intervention, we utilized a 3D cerebral organoid model derived from human pluripotent stem cells generated from an AD patient. We treated cerebral organoids carrying ApoE3 and ApoE4 variants with beta and gamma-secretase inhibitors to determine if organoids could reproduce the differences observed in the 2D model and if the alteration in secretase activity could affect the regulation of neuronal lipids synthesis in an ApoE-dependent manner. Ganglioside profiling was accomplished using liquid chromatography/electrospray ionization tandem mass spectrometry (LC/ESI-MS/MS) via selective reaction monitoring (SRM). The inhibitor administration elevated the levels of ganglioside and ceramide in ApoE4-derived organoids. Since gangliosides are known to enhance Aβ fibrillogenesis, our study implies that reduction in secretase activity affects neuronal membrane architecture that could eventually aggravate AD, particularly in patients with the ApoE4 isoform. In addition, the ability of organoids to replicate results from other experimental models demonstrates their potential to improve translatability significantly.

## Introduction

Alzheimer’s disease (AD) is a neurodegenerative disorder marked by a progressive loss of the structure and function of neurons, leading to cell death. This could result in the deterioration of cognitive and motor skills. It is the most frequent cause of dementia and typically affects those over 65^1^. Although the precise etiology of AD is unknown, a mix of malfunctions in complex regulatory processes, such as endocytosis, lipid metabolism, and vesicle trafficking, have been identified^2–4^.

One of the hallmarks of AD is the formation of amyloid beta (Aβ) plaques in the brain. Beta and gamma secretases cleave amyloid precursor protein (APP) and contribute to forming Aβ plaques. This makes secretase inhibitors a fascinating pharmacological target for lowering Aβ synthesis and slowing the course of AD^5^. It is well recognized that secretases, such as β-site APP cleaving enzyme 1 (BACE1), considerably impact neuronal lipid metabolism and function^6,7^. Lipids are abundant in the brain and essential for sustaining normal brain processes such as supporting the structural integrity of the brain, enabling signal transduction, and myelination^4^. Disruption in lipid metabolism is linked to various neurological diseases, including AD^8^. This could posit that disturbances in secretase activity may lead to changed lipids levels, thus increasing the risk for AD^9,10^. Sialylated glycosphingolipids called gangliosides are widely distributed throughout the brain, particularly in the neurons. Studies have reported that BACE1 could regulate ganglioside metabolism by affecting the activity of other enzymes involved in their synthesis and degradation, potentially modulating their levels^6^. BACE1 has also been shown to cleave gangliosides and generate new ganglioside species that may have different biological activities than the original ^11^. The precise mechanisms by which secretases affect gangliosides are not fully understood and require further research.

The pathogenesis of AD is complex, encompassing a wide array of cell types, tissue structures, and metabolic pathways. Additionally, the absence of an adequate disease model that accurately reproduces the complexity and pathological features of this human-specific disease, coupled with the difficulty of acquiring human brain samples, presents a significant challenge. Nevertheless, recent advancements in human pluripotent stem cells-derived cerebral organoids, three-dimensional representations of the human brain ^12^, have provided new insights into AD ^13^. The organoids have demonstrated relevant features of AD, such as the production of amyloidogenic variants of the Aβ protein and increased tau phosphorylation, as well as the ability to form a variety of brain-like regions with complex tissues^14,15^. Furthermore, existing animal transgenic models fall short in replicating sporadic AD (sAD), which is the most prevalent form of the disease. Notably, the presence of the ApoE4 allele significantly increases the risk of sAD^16^. To address this limitation, organoids can play a pivotal role by incorporating specific genetic mutations for more accurate modeling and study of sAD ^17^.

In this study, we investigated the impact of inhibiting β and γ-secretase activity on the production of Aβ in patients’ derived cerebral organoids. We treated organoids with secretase inhibitors and found a significant decrease in Aβ40 and Aβ42 levels, with no difference between ApoE genotypes. We also assessed the effect of these inhibitors on neuronal lipids. Due to increased sensitivity, specificity, and precision, a mass spectrometry (MS) assay via selected reaction monitoring (SRM) was employed to analyze gangliosides. In addition to quantifying major gangliosides, the assay was also utilized to identify the molecular composition of lipids, including the fatty acid composition and the head group structure. We identified 70 molecular species of gangliosides in a single organoid, with C16 and C18 fatty acyl chains being the most prevalent. Importantly, suppressing secretase activity increased ganglioside levels in sAD-E4 organoids but not in sAD-E3 organoids. Lastly, the exogenous addition of GM1 to cerebral organoids showed that increased GM1 level does not lead to the accumulation of APP clusters associated with Alzheimer’s disease pathology. In sum, by employing our MS-based ganglioside profiling assay in the brain organoids, we identified a hitherto unreported link in which ApoE allele-dependent ganglioside metabolism dysregulation was observed upon secretase inhibition in cerebral organoids.

## Materials and Methods

### Materials

The BACE1 inhibitor was purchased as BACE inhibitor C3 (β-Secretase Inhibitor IV), and the gamma secretases inhibitor was purchased as Compound E (γ-Secretase Inhibitor XXI) from EMD Millipore (Burlington, MA, USA). Ammonium acetate (Cat# A11450) and ammonium formate (Cat# A11550) were from Fisher Chemical (Hampton, NH, USA). LC-MS grade acetonitrile (ACN, Cat# 34967), isopropanol (IPA, Cat# 34965), and formic acid (FA, for MS ∼98% purity) were from Honeywell (Charlotte, NC, USA). LC-MS grade methanol (Cat# 0013687802BS) was from Biosolve Chimie (Dieuze, France). Acetic acid (≥99.8%) was from Penta Chemicals (Chrudim, Czech Republic). Isotopically-labeled ganglioside internal standards (GM1 and GM3) were synthesized by Lukas Opalka (Faculty of Pharmacy, Charles University, Prague, Czech Republic). The ultrapure water was prepared in the purification system (arium® Comfort System, Sartorius, Gottingen, Germany).

### Cerebral organoid culture and treatment with β and γ-secretase inhibitors

For the generation of 3D cultures of cerebral organoids, we used organoids derived from five independent human induced pluripotent stem cell (iPSC) lines. Specifically, we used one pair of isogenic iPSCs where the original iPSC line was derived from an sAD patient carrying the ApoE4/4 genotype, and we refer to this line as “sAD-E4”. This cell line was then edited using CRISPR/Cas9, and instead of the ApoE4/4 genotype, it now carries the ApoE3/3 gene variant. We refer to this iPSC line as “sAD-E3”. This set of iPSC lines was generously provided by Dr. Li-Huei Tsai and previously described^18^. For experiments with exogenously added GM1 and/or subsequent APP-cluster formation assessment, we used three iPSC lines derived from two healthy non-demented individuals carrying ApoE3/3 genotype (MUNIi008-A, MUNIi010-A, here referred to as “ApoE3/3”) and one healthy non-demented individual carrying ApoE3/4 genotype (MUNIi009-A, here referred to as “ApoE3/4”). These cell lines were derived from fibroblasts deposited at the Coriell Cell bank repository, registered at https://hpscreg.eu/, and characterized^19^. All iPSC lines were ^1^maintained under standard culture conditions as previously described^20^, using mTESR medium (Stem Cell Technologies) on cell culture plates coated with Geltrex (Thermo Fisher Scientific). Organoids for the experiments described in this study were generated following previously published protocols^12,15,20,21^. To treat the organoids with β and γ-Secretase inhibitors, we followed a previously published protocol with minor modifications^13^. Specifically, 50-day-old organoids derived from both isogenic iPSCs were treated for 50 days with media change every other day. BACE Inhibitor C3 (5 μM) and Compound E (6nM) were added to the differentiation medium freshly. Organoids were harvested from two independent batches of CO differentiation (N=5 each). As control samples, we used non-treated (NTR) organoids or organoids treated with dimethyl sulfoxide (DMSO), the solvent used to dilute the inhibitors.

### The enzyme-linked immunosorbent assay (ELISA)

For the ELISA, individual cerebral organoids (COs) were cultured in Essential 6™ Medium (Thermo Fisher Scientific) for 72 hours before undergoing analysis. Subsequently, the culture medium and the corresponding organoid were harvested and kept separately at -80°C. The quantification of Aβ40 and Aβ42 peptides present in the cell culture medium was performed utilizing the Amyloid beta 40 Human ELISA Kit (Thermo Fisher Scientific) and the Amyloid beta 42 Human ELISA Kit, Ultrasensitive (Thermo Fisher Scientific), following the instructions provided by the manufacturer. Samples were analyzed in technical duplicates to ensure accuracy. To compare the Aβ40 and Aβ42 peptide levels across organoids of varying sizes, we determined the total protein concentration in lysed organoids, computed the total protein weight, and employed this value for normalization purposes. The resulting measurements are expressed in picograms of Aβ per microgram of protein.

### Treatment of organoids with Monosialoganglioside (GM1)

To test the incorporation of GM1 to the cell membranes of differentiating cells within cerebral organoids, we used fluorescently labeled GM1-FL-BODIPY, generously provided by Dr. Martin Hof and Dr. Radek Sachl, Heyrovsky Institute, Prague. GM1-FL-BODIPY was added at the 50 μM concentration directly to the cell culture medium and 50-day-old organoids were grown in this medium for 3 days. Subsequently, organoids were washed and fixed using 3.7% paraformaldehyde (PFA) for one hour. Intact organoids were then stored in PBS at +4°C until further processing for immunohistochemistry.

For experiments with short-term and long-term treatment of organoids with GM1, we used commercially available, non-fluorescent GM1 at the concentration 50 μM, which was freshly added to the cell culture medium for every medium change. We initiated the exposure of organoids at day 60 and continued the treatment until Day 135 for a total of 65 days. Organoids were then washed in PBS, fixed in 3.7% PFA, and stored in PBS at +4°C until further processing for immunohistochemistry.

### Immunohistochemistry and evaluation of APP accumulation

Before the immunostaining process, the collected cerebral organoids were subjected to fixation using a 3.7% PFA solution for one hour.

For cryo-sections, fixed and stored organoids were saturated with 30% sucrose (Merck), embedded in O.C.T. medium (Tissue-Tek), and frozen. Subsequently, 10 μm sections were prepared on cryostat Leica 1850. Excessing O.C.T. medium was removed by a 15-minute PBS wash prior to immunohistochemistry staining.

For the immunohistochemistry (IHC) procedure, histological crysections of the cerebral organoids were initially permeabilized using a 0.2% Triton-X solution (Merck) in phosphate-buffered saline (PBS) and subsequently blocked in a solution containing 2% normal goat serum (Merck) within the same permeabilization solution. The sections were then incubated overnight at 4°C with an Anti-APP primary antibody (D54D2, Cell Signaling), diluted in the blocking solution. Subsequently, a secondary antibody was applied for a one-hour incubation at room temperature. Nuclei were visualized using 4’,6-diamidino-2-phenylindole (DAPI; Carl Roth). The histological sections were imaged utilizing an inverted microscope Zeiss Axio Observer.Z1 equipped with a confocal unit LSM 800 (Zeiss).

For the quantification of β-amyloid (Aβ) aggregates, paraffin sections were employed for IHC and then scanned using the fluorescent microscope TissueFAXS (TissueGnostics GmbH). To quantify these aggregates, a minimum of four images were analyzed for each organoid section. For each cell line and treatment, at least four separate sections from different regions were evaluated, and two organoids were sectioned per time point. The size of Aβ aggregates was determined through analysis using ImageJ software. To establish a threshold and eliminate the IHC background, the threshold value was set at 20 pixels. Aβ aggregates were identified as APP clusters exceeding 100 pixels in size. The percentage of larger Aβ aggregates was calculated for each section and represented as individual data points on a graph. These values were then normalized relative to the respective control averages.

### Sample preparation and Liquid chromatography-electrospray ionization-tandem mass spectrometry (LC/ESI-MS/MS) for the analysis of gangliosides

Lipid extraction followed by sequential protein extraction was performed, as previously reported by our group^21^. Briefly, the harvested organoids were freeze-dried and homogenized. Lipids were extracted upon the addition of 80% IPA to the dry homogenate, followed by vortex and sonication. The sample was further centrifuged, and the lipid extract was removed from the residual protein pellet. The lipid extracts were stored at - 20°C until analysis.

The lipid extract was mixed with isotopically labeled GM1 (0.3 μM) and GM3 (0.15 μM) internal standards for relative ganglioside quantitation. For the ganglioside assay, ApoE3/3 (N=10) and ApoE4/4 (N=10) control organoids, along with ApoE3/3 (N=10) and ApoE4/4 (N=10) sAD organoids were utilized. The analysis of major species of gangliosides through LC/ESI-MS/MS was followed based on the same protocol^21^. Using an Agilent 1200 HPLC system coupled to an ESI 6495 Triple Q MS, we analyzed eight subclasses of gangliosides – GD3, GM3, GM2, GD2, GM1, GD1a, GD1b, and GT1b. We also analyzed molecular species with different ceramide chains in each subclass. They were characterized using their retention time and the total carbon number of the lipids-which is manifested in gangliosides through their fatty acid chain and degree of saturation^22^. A detailed list of the SRM transitions is available in Table S1. Data were processed and analyzed in MassHunter Quantitative Analysis software (Agilent Technologies, Santa Clara, USA).

### Statistical analysis

The graphs were plotted using GraphPad Prism 9.0.0 (GraphPad Software, Inc., La Jolla, CA, USA). The data were normally distributed, as assessed by the Shapiro-Wilk test (p > 0.05). Groups were compared using Student’s two-tailed t-test (two groups). To identify any separation in the widespread data, principal component analysis was performed using an open-access tool on Metaboanalyst (Metaboanalyst 5.0^23^). The data were scaled using the z-score method before PCA. The figures were prepared using BioRender scientific illustration software (BioRender.com). Details are provided in individual figure legends.

## Results

### 1. The effect of suppression of secretase activity on Aβ production in cerebral organoids

First, to determine the effect of inhibitors on the levels of secreted amyloid β peptides, we treated sAD-E4 and sAD-E3 (N=4) cerebral organoids (N=4) with β and γ-secretase inhibitors and measured the levels of Aβ40 and Aβ42 production to the cell culture media by ELISA. We used NTR and DMSO-treated organoids (N=4) as controls. Our results show that upon inhibition of both β and γ-secretase, there was a significant decrease in Aβ40 and Aβ42 levels in both sAD-E4 and sAD-E3 organoids (Fig 1). No differential change was observed between E3 and E4 organoids upon inhibitor treatment. As expected, we also did not observe any significant difference in the Aβ40 and Aβ42 levels between NTR and DMSO conditions, irrespective of the ApoE genotype. Data thus suggest that organoids adequately respond to the treatment by secretase inhibitors and that this experimental setup could thus be used for all our subsequent analyses.

**Fig 1.**
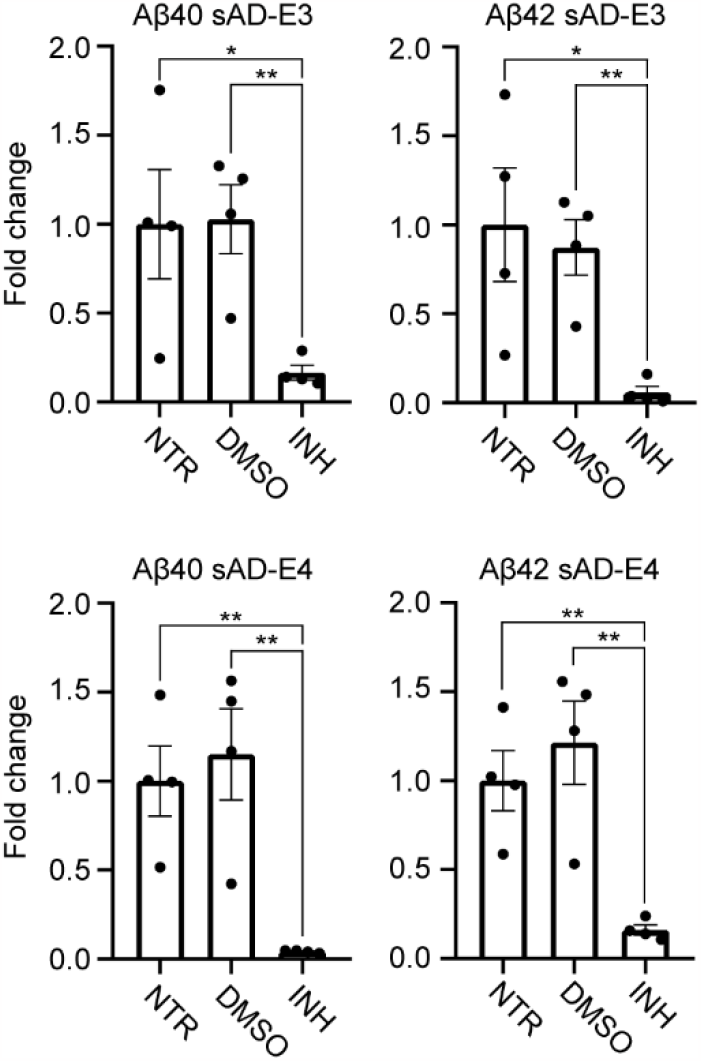
Effect of beta and gamma-secretase inhibitor activity on Aβ production. sAD-E4 and sAD-E3 organoids were treated with BACE inhibitor C3 and Compound E (INH) for 50 days, and levels of Aβ40 and Aβ42 in the cell culture media were determined by ELISA. Each dot represents a measurement of cell culture media concentration from a single organoid (N=4). The levels of DMSO and INH are relative to the non-treated (NTR) levels.

### 2. Characterization of gangliosides in the cerebral organoids

Separation and classification of gangliosides according to their ceramide type was done in the RPLC column in both positive and negative ionization mode. To validate the method, we used organoids derived from both sAD-E3 and sAD-E4 cell lines. Predictably, gangliosides were separated from the most hydrophilic to the hydrophobic molecules. Precursor ions with GM1, GM2, and GM3 were detected in the state of a single charge, with GD1 and GT1 detected in the state of a double charge. A common fragment ion at m/z 290 composed of sialic acid ions (SA2) was used as the product ion in the negative mode. The ceramide moiety of gangliosides influenced the elution of different molecular species. The retention behavior of gangliosides followed the patterns related to the number of carbon atoms in fatty acyl chains and the number of double bonds (Fig 2). Gangliosides were eluted from d16:1-18:0 (d18:1-16:0) to d18:1-22:0. We could also distinguish between regioisomeric gangliosides such as GD1a and GD1b using product ions such as SA-SA2 (/m/z 581). In total, we profiled eight subclasses of gangliosides, identifying 70 molecular species in a single cerebral organoid. Interestingly, the most prevalent species were C16 and C18 fatty acyl chains, in contrast to the prevalent C18 and C20 fatty acyl chains in adult human brains^24^.

**Fig 2.**
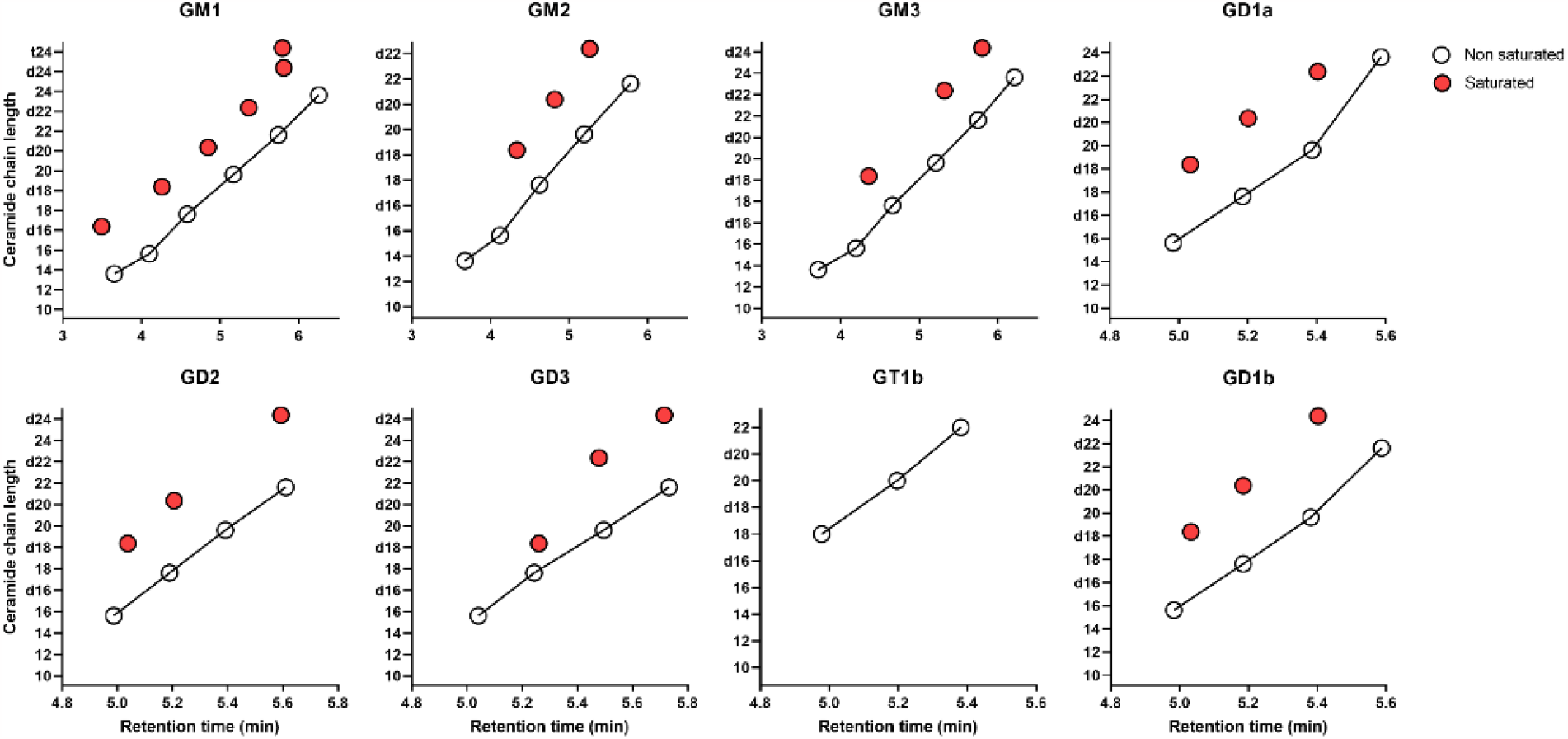
The retention time of gangliosides profiled in cerebral organoids for fatty acid chain lengths ranging from C14:0 to C24:0. The red dots (o) indicate the non-saturated fatty acid chains, whereas the white dots (o) indicate saturated fatty acid chains. A total of 60 molecular species were characterized in a single cerebral organoid.

### 3. Suppression of secretase activity increased the levels of gangliosides in an ApoE-dependent manner in cerebral organoids

Subsequently, we aimed to characterize the composition of gangliosides in sAD-E4 and sAD-E3 organoids. Thus, based on our method to characterize gangliosides (as described in the previous section), we performed relative quantitation of the species mentioned above in two batches of organoids (N=10).

PCA plot of the organoids showed no separation in the ApoE3/3 isoform upon treatment with β and γ-Secretase inhibitors; however, we observed slight separation in the ApoE4/4 line (Fig 3a). Upon closer look at the total ganglioside concentrations, we observed a significant increase in their levels in sAD-E4 organoids treated with inhibitors. These results were limited to the ApoE4/4 line alone, with no upregulation observed in the ApoE3/3 line (Fig 3b, Fig S1).

**Fig 3.**
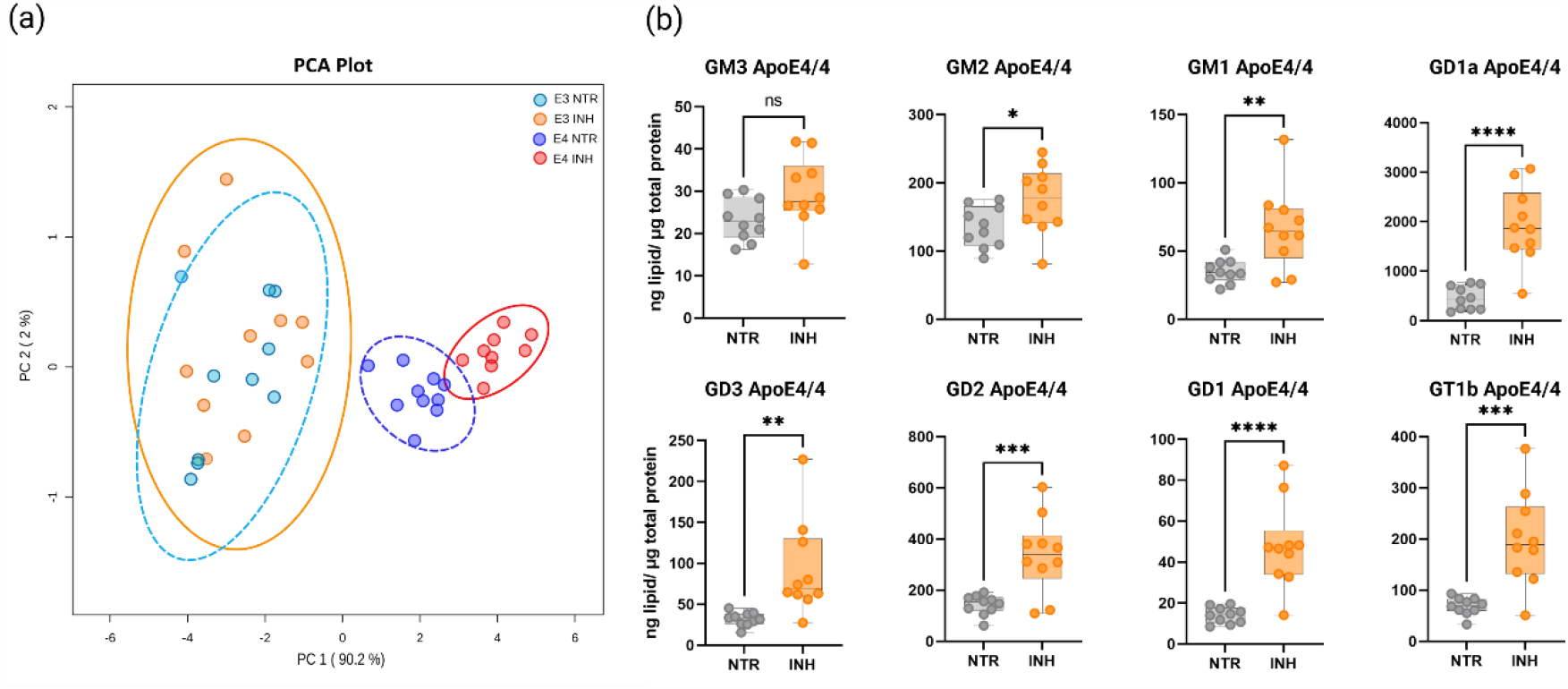
Effect of reduction of secretase activity on levels of gangliosides. sAD-E3 and sAD-E4 organoids were treated with BACE inhibitor C3 and Compound E (INH) for 50 days. The level of gangliosides in the NTR and inhibitor-treated organoids were measured by LC-ESI/MS in combination with SRM. (a) Principal component analysis (PCA) of N=10 organoids, each from NTR sAD-E3, NTR sAD-E4, INH sAD-E3, and INH sAD-E4 conditions. (b) Total ganglioside levels in NTR and INH cerebral organoids. Each box plot indicates the minimum and maximum with a median of 10 values. ****P < 0.0001, ***P < 0.001, **P < 0.01, *P < 0.05 (Groups were compared using two-tailed t-test).

We were also interested in the effect of inhibitors on the precursors involved in the formation of gangliosides. Strikingly, we also found an increase in dihydroceramide and ceramide species in the sAD-E4 organoids, which contrasted with the sAD-E3 organoids, where their levels had significantly declined upon inhibitor treatment (Fig S1 and S2).

### 4. Exogenous addition of GM1 to ApoE3 and ApoE4 genotype cerebral organoids

Lastly, we aimed to verify whether perturbation in the GM1 level could influence the development of Alzheimer’s disease-like pathology, i.e., accumulation of APP plaques. We thus first tested if the exogenously added GM1 directly to the cell culture medium would be incorporated into the cerebral organoids and metabolized by them. Results show that our fluorescently labeled GM1 can be detected inside the cerebral organoids, indicating that organoids indeed uptake this ganglioside (Fig 4a). Subsequent MS analysis confirmed that organoids treated with GM1 not only increased the level of GM1 but also increased levels of GM2 and GD1a species, which are direct degradation products and precursors of GM1, respectively. This data indicated the active metabolism of GM1 inside the cells (Fig 4b).

**Fig 4.**
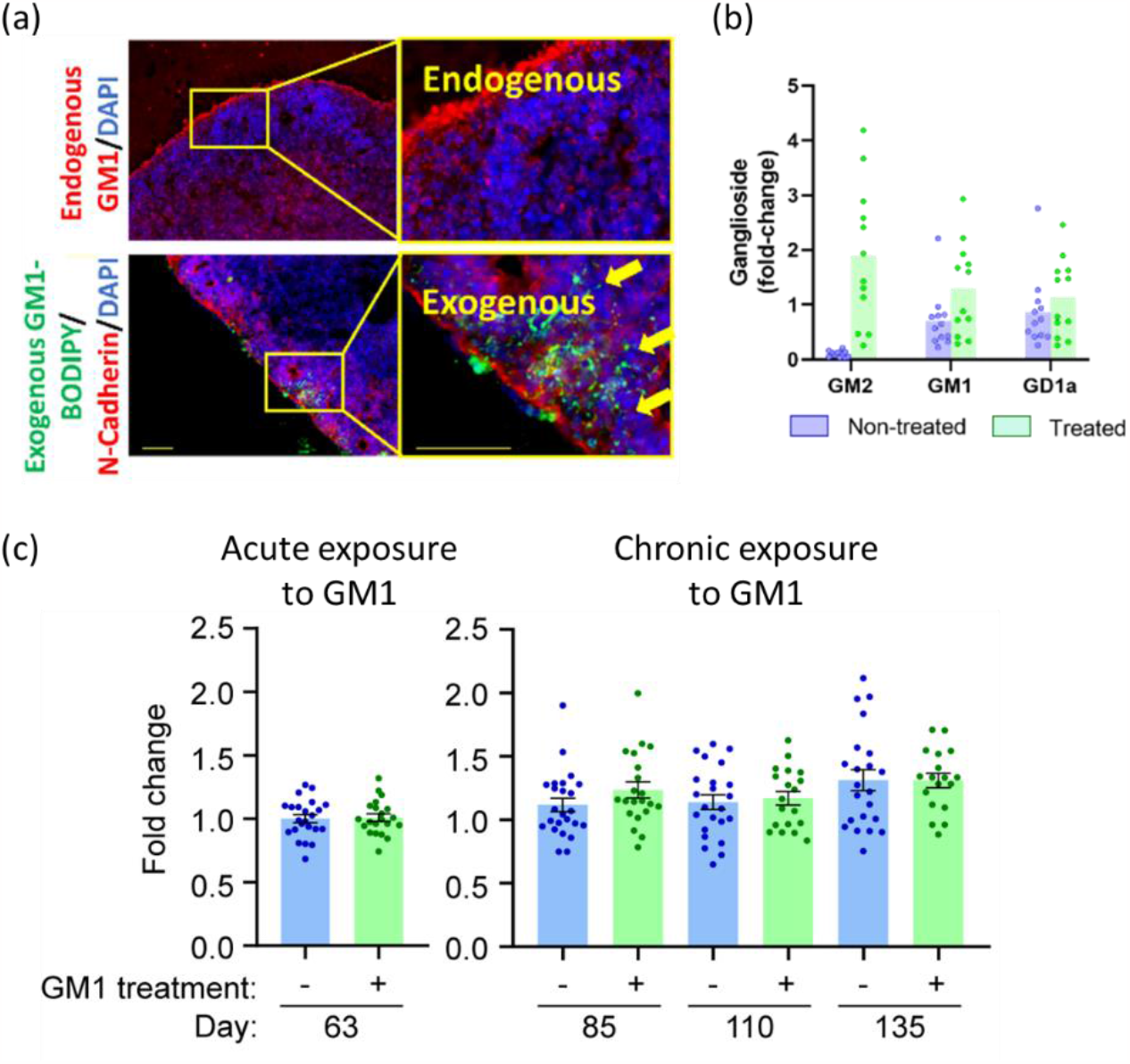
Effect of application of exogenous GM1 on Aβ and ganglioside production. (a) Indirect immunofluorescent staining images depict the endogenous GM1 and the labeled exogenous GM1 in the cerebral organoids. (b) Total ganglioside levels of GM1, GM2, and GD1a in NTR and GM1 treated organoids in ApoE3/3 organoids. The columns represent the average ±standard deviation. The dots represent individual organoids. (c) The accumulation of APP clusters evaluated from IHC analysis of APOE3/3 and APOE3/4 organoids after Acute (3 days) and Chronic (25, 50, and 75 days) exposure to GM1.

Finally, we performed experiments where we exogenously added GM1 to 60-day-old organoids for three days (Acute exposure) as well as during long-term cell culture for 75 days (Chronic exposure). During this long-term exposure, we also harvested several time points to map the progress of possible APP cluster formation. Results show that despite increased levels of GM1 during Acute and Chronic exposure, cerebral organoids did not accumulate APP clusters (Fig 4c). These findings suggest that an increased GM1 level in the cell membrane does not directly induce the accumulation of APP or the formation of APP clusters, characteristics of Alzheimer’s disease pathology.

### Discussion

As anticipated, treating cerebral organoids with secretase inhibitors led to a considerable decrease in Aβ40 and Aβ42 relative to control organoids in the E3 and E4 lines. Interestingly, we also detected a minor rise in Aβ40 levels relative to Aβ42 in the ApoE3 line, in sharp contrast to the ApoE4/4 line, which had greater Aβ42 levels than Aβ40 levels (Fig 1). A plausible explanation could be ApoE4’s ability to bind to the secretase complex and suppress the synthesis of Aβ40, as previously reported^25^.

There is evidence that the accumulation of gangliosides, GM3 and GM1, can facilitate the development of the GAβ complex, resulting in a conformational shift serving as a seed for fibrillogenesis^26–29^. An age-dependent rise in GM1 has also been found in synaptosomes containing the ApoE4 isoform, but the increase was not as significant in synaptosomes containing the ApoE3 isoform^30^. We utilized a targeted approach established for studying gangliosides to profile molecular species relative to one another. Thus, the accumulation of gangliosides observed in the sAD-E4 cerebral organoids indicates the potential to aggravate AD etiology further.

Secretases such as BACE1 are known to be elevated in the brains of AD patients, making them an attractive therapeutic target^16,31^. Elevated expression of GM1, GM2, or GD2 increases β- and γ-site cleavages of APP^32^. β-1,4 N-acetylgalactosaminyltransferase (B4GALNT1), one of the glycosyltransferases responsible for producing these complex gangliosides, is known to affect BACE1 stability by inhibiting its lysosomal degradation and boosting its protein levels^6^. This ultimately results in a surge of the beta-cleavage of APP. The ApoE4 isoform, a known hereditary risk factor for AD, can not only exacerbate neurodegeneration^16,33^ but can also influence the activity of γ -secretase by raising the amount of γ-secretase complex attached to it, thus escalating the synthesis of Aβ42^34^. This demonstrates an unsolved mechanism through which ApoE isoform-dependent neuronal lipid dysregulation mediated by decreased secretase activity may contribute to AD pathogenesis. Upregulation of ganglioside metabolism previously observed in neuritic cell lines treated with secretase inhibitors was replicated precisely in organoids^9^ (Fig 3b).

Our attempts to detect dysregulation in Aβ levels upon organoid treatment with exogenous GM1 were unsuccessful (Fig 4b). It is conceivable that the mere incorporation of this lipid into the cell surface probably does not affect APP processing. As previously documented, surface-adherent exogenous gangliosides cannot alter the stability of BACE1, resulting in no change in APP cleavage^6^.

## Conclusion

In conclusion, we demonstrated that a decrease in secretase activity modifies ganglioside levels in ApoE4 genotype cerebral organoids, which could contribute to AD pathology’s genesis. In the future, it will be intriguing to see if ApoE4’s propensity to sequester secretases directly impacts glycosyltransferases and ganglioside levels. Modulation of ganglioside metabolism could present an opportunity to develop a way to prevent and treat AD. Our study further proved the efficacy of cerebral organoids in exhibiting AD-like pathophysiology. This model has the potential to dramatically expedite research and drug discovery in Alzheimer’s disease, as well as provide valuable insights into its underlying causes.

## Author Information

^*^ Correspondence should be addressed to Dr. Zdenek Spacil Phone: (+420) 549 49 7989.

## References

(1) 2022 Alzheimer’s Disease Facts and Figures. Alzheimer’s & Dementia 2022, 18 (4), 700–789. 10.1002/alz.12638.

(2) Tang, B. L. Neuronal Protein Trafficking Associated with Alzheimer Disease. Cell Adh Migr 2009, 3 (1), 118–128. 10.4161/cam.3.1.7254.

(3) Domínguez-Prieto, M.; Velasco, A.; Tabernero, A.; Medina, J. M. Endocytosis and Transcytosis of Amyloid-β Peptides by Astrocytes: A Possible Mechanism for Amyloid-β Clearance in Alzheimer’s Disease. Journal of Alzheimer’s Disease 2018, 65 (4), 1109–1124. 10.3233/JAD-180332.

(4) Chew, H.; Solomon, V. A.; Fonteh, A. N. Involvement of Lipids in Alzheimer’s Disease Pathology and Potential Therapies. Front Physiol 2020, 11, 598. 10.3389/fphys.2020.00598.

(5) Loeffler, D. A. Experimental Approaches for Altering the Expression of Abeta-degrading Enzymes. J Neurochem 2023. 10.1111/jnc.15762.

(6) Yamaguchi, T.; Yamauchi, Y.; Furukawa, K.; Ohmi, Y.; Ohkawa, Y.; Zhang, Q.; Okajima, T.; Furukawa, K. Expression of B4GALNT1, an Essential Glycosyltransferase for the Synthesis of Complex Gangliosides, Suppresses BACE1 Degradation and Modulates APP Processing. Sci Rep 2016, 6 (1), 34505. 10.1038/srep34505.

(7) Takasugi, N.; Sasaki, T.; Suzuki, K.; Osawa, S.; Isshiki, H.; Hori, Y.; Shimada, N.; Higo, T.; Yokoshima, S.; Fukuyama, T.; Lee, V. M.-Y.; Trojanowski, J. Q.; Tomita, T.; Iwatsubo, T. BACE1 Activity Is Modulated by Cell-Associated Sphingosine-1-Phosphate. Journal of Neuroscience 2011, 31 (18), 6850–6857. 10.1523/JNEUROSCI.6467-10.2011.

(8) Arbor, S. C.; Lafontaine, M.; Cumbay, M. Amyloid-Beta Alzheimer Targets — Protein Processing, Lipid Rafts, and Amyloid-Beta Pores. Yale Journal of Biology and Medicine. Yale Journal of Biology and Medicine Inc. March 1, 2016, pp 5–21. /pmc/articles/PMC4797837/ (accessed 2021-02-10).

(9) Oikawa, N.; Goto, M.; Ikeda, K.; Taguchi, R.; Yanagisawa, K. The γ-Secretase Inhibitor DAPT Increases the Levels of Gangliosides at Neuritic Terminals of Differentiating PC12 Cells. Neurosci Lett 2012, 525 (1), 49–53. 10.1016/j.neulet.2012.07.027.

(10) Holmes, O.; Paturi, S.; Ye, W.; Wolfe, M. S.; Selkoe, D. J. Effects of Membrane Lipids on the Activity and Processivity of Purified γ-Secretase. Biochemistry 2012, 51 (17), 3565–3575. 10.1021/bi300303g.

(11) Moll, T.; Shaw, P. J.; Cooper-Knock, J. Disrupted Glycosylation of Lipids and Proteins Is a Cause of Neurodegeneration. Brain 2020, 143 (5), 1332–1340. 10.1093/brain/awz358.

(12) Lancaster, M. A.; Knoblich, J. A. Generation of Cerebral Organoids from Human Pluripotent Stem Cells. Nat Protoc 2014, 9 (10), 2329–2340. 10.1038/nprot.2014.158.

(13) Raja, W. K.; Mungenast, A. E.; Lin, Y.-T.; Ko, T.; Abdurrob, F.; Seo, J.; Tsai, L.-H. Self-Organizing 3D Human Neural Tissue Derived from Induced Pluripotent Stem Cells Recapitulate Alzheimer’s Disease Phenotypes. PLoS One 2016, 11 (9), e0161969. 10.1371/journal.pone.0161969.

(14) Gonzalez, C.; Armijo, E.; Bravo-Alegria, J.; Becerra-Calixto, A.; Mays, C. E.; Soto, C. Modeling Amyloid Beta and Tau Pathology in Human Cerebral Organoids. Mol Psychiatry 2018, 23 (12), 2363–2374. 10.1038/s41380-018-0229-8.

(15) Camp, J. G.; Badsha, F.; Florio, M.; Kanton, S.; Gerber, T.; Wilsch-Bräuninger, M.; Lewitus, E.; Sykes, A.; Hevers, W.; Lancaster, M.; Knoblich, J. A.; Lachmann, R.; Pääbo, S.; Huttner, W. B.; Treutlein, B. Human Cerebral Organoids Recapitulate Gene Expression Programs of Fetal Neocortex Development. Proceedings of the National Academy of Sciences 2015, 112 (51), 15672–15677. 10.1073/pnas.1520760112.

(16) Zhao, J.; Fu, Y.; Yamazaki, Y.; Ren, Y.; Davis, M. D.; Liu, C.-C.; Lu, W.; Wang, X.; Chen, K.; Cherukuri, Y.; Jia, L.; Martens, Y. A.; Job, L.; Shue, F.; Nguyen, T. T.; Younkin, S. G.; Graff-Radford, N. R.; Wszolek, Z. K.; Brafman, D. A.; Asmann, Y. W.; Ertekin-Taner, N.; Kanekiyo, T.; Bu, G. APOE4 Exacerbates Synapse Loss and Neurodegeneration in Alzheimer’s Disease Patient IPSC-Derived Cerebral Organoids. Nat Commun 2020, 11 (1), 5540. 10.1038/s41467-020-19264-0.

(17) Park, J.-C.; Jang, S.-Y.; Lee, D.; Lee, J.; Kang, U.; Chang, H.; Kim, H. J.; Han, S.-H.; Seo, J.; Choi, M.; Lee, D. Y.; Byun, M. S.; Yi, D.; Cho, K.-H.; Mook-Jung, I. A Logical Network-Based Drug-Screening Platform for Alzheimer’s Disease Representing Pathological Features of Human Brain Organoids. Nat Commun 2021, 12 (1), 280. 10.1038/s41467-020-20440-5.

(18) Lin, Y. T.; Seo, J.; Gao, F.; Feldman, H. M.; Wen, H. L.; Penney, J.; Cam, H. P.; Gjoneska, E.; Raja, W. K.; Cheng, J.; Rueda, R.; Kritskiy, O.; Abdurrob, F.; Peng, Z.; Milo, B.; Yu, C. J.; Elmsaouri, S.; Dey, D.; Ko, T.; Yankner, B. A.; Tsai, L. H. APOE4 Causes Widespread Molecular and Cellular Alterations Associated with Alzheimer’s Disease Phenotypes in Human IPSC-Derived Brain Cell Types. Neuron 2018, 98 (6), 1141-1154.e7. 10.1016/j.neuron.2018.05.008.

(19) Raska, J.; Hribkova, H.; Klimova, H.; Fedorova, V.; Barak, M.; Barta, T.; Pospisilova, V.; Vochyanova, S.; Vanova, T.; Bohaciakova, D. Generation of Six Human IPSC Lines from Patients with a Familial Alzheimer’s Disease (n = 3) and Sex- and Age-Matched Healthy Controls (n = 3). Stem Cell Res 2021, 53. 10.1016/j.scr.2021.102379.

(20) Nemergut, M.; Marques, S. M.; Uhrik, L.; Vanova, T.; Nezvedova, M.; Gadara, D. C.; Jha, D.; Tulis, J.; Novakova, V.; Planas-Iglesias, J.; Kunka, A.; Legrand, A.; Hribkova, H.; Pospisilova, V.; Sedmik, J.; Raska, J.; Prokop, Z.; Damborsky, J.; Bohaciakova, D.; Spacil, Z.; Hernychova, L.; Bednar, D.; Marek, M. Domino-like Effect of C112R Mutation on ApoE4 Aggregation and Its Reduction by Alzheimer’s Disease Drug Candidate. Mol Neurodegener 2023, 18 (1). 10.1186/s13024-023-00620-9.

(21) Nezvedová, M.; Jha, D.; Váňová, T.; Gadara, D.; Klímová, H.; Raška, J.; Opálka, L.; Bohačiaková, D.; Spáčil, Z. Single Cerebral Organoid Mass Spectrometry of Cell-Specific Protein and Glycosphingolipid Traits. Anal Chem 2023. 10.1021/ACS.ANALCHEM.2C00981.

(22) Cajka, T.; Fiehn, O. Comprehensive Analysis of Lipids in Biological Systems by Liquid Chromatography-Mass Spectrometry. TrAC - Trends in Analytical Chemistry. Elsevier B.V. October 1, 2014, pp 192–206. 10.1016/j.trac.2014.04.017.

(23) Xia, J.; Psychogios, N.; Young, N.; Wishart, D. S. MetaboAnalyst: A Web Server for Metabolomic Data Analysis and Interpretation. Nucleic Acids Res 2009, 37 (SUPPL. 2). 10.1093/nar/gkp356.

(24) Caughlin, S.; Maheshwari, S.; Weishaupt, N.; Yeung, K. K. C.; Cechetto, D. F.; Whitehead, S. N. Age-Dependent and Regional Heterogeneity in the Long-Chain Base of A-Series Gangliosides Observed in the Rat Brain Using MALDI Imaging. Sci Rep 2017, 7 (1), 1–12. 10.1038/s41598-017-16389-z.

(25) Sun, Y.; Islam, S.; Gao, Y.; Nakamura, T.; Zou, K.; Michikawa, M. Apolipoprotein E4 Inhibits γ-Secretase Activity via Binding to the γ-Secretase Complex. J Neurochem 2023. 10.1111/JNC.15750.

(26) Yanagisawa, K.; Odaka, A.; Suzuki, N.; Ihara, Y. GM1 Ganglioside–Bound Amyloid β–Protein (Aβ): A Possible Form of Preamyloid in Alzheimer’s Disease. Nat Med 1995, 1 (10), 1062–1066. 10.1038/nm1095-1062.

(27) Peters, I.; Igbavboa, U.; Schütt, T.; Haidari, S.; Hartig, U.; Rosello, X.; Böttner, S.; Copanaki, E.; Deller, T.; Kögel, D.; Wood, W. G.; Müller, W. E.; Eckert, G. P. The Interaction of Beta-Amyloid Protein with Cellular Membranes Stimulates Its Own Production. Biochimica et Biophysica Acta (BBA) - Biomembranes 2009, 1788 (5), 964–972. 10.1016/j.bbamem.2009.01.012.

(28) Grimm, M. O. W.; Zinser, E. G.; Grösgen, S.; Hundsdörfer, B.; Rothhaar, T. L.; Burg, V. K.; Kaestner, L.; Bayer, T. A.; Lipp, P.; Müller, U.; Grimm, H. S.; Hartmann, T. Amyloid Precursor Protein (APP) Mediated Regulation of Ganglioside Homeostasis Linking Alzheimer’s Disease Pathology with Ganglioside Metabolism. PLoS One 2012, 7 (3), e34095. 10.1371/journal.pone.0034095.

(29) Cebecauer, M.; Hof, M.; Amaro, M. Impact of GM1 on Membrane-Mediated Aggregation/Oligomerization of β-Amyloid: Unifying View. Biophys J 2017, 113 (6), 1194–1199. 10.1016/j.bpj.2017.03.009.

(30) Yamamoto, N.; Igbabvoa, U.; Shimada, Y.; Ohno-Iwashita, Y.; Kobayashi, M.; Wood, W. G.; Fujita, S. C.; Yanagisawa, K. Accelerated Aβ Aggregation in the Presence of GM1-Ganglioside-Accumulated Synaptosomes of Aged ApoE4-Knock-in Mouse Brain. FEBS Lett 2004, 569 (1–3), 135–139. 10.1016/j.febslet.2004.05.037.

(31) Imbimbo, B. P.; Ippati, S.; Watling, M.; Imbimbo, C. Role of Monomeric Amyloid-β in Cognitive Performance in Alzheimer’s Disease: Insights from Clinical Trials with Secretase Inhibitors and Monoclonal Antibodies. Pharmacol Res 2023, 187 (October 2022), 106631. 10.1016/j.phrs.2022.106631.

(32) Giraudo, C. G.; Rosales Fritz, V. M.; Maccioni, H. J. F. GA2/GM2/GD2 Synthase Localizes to the Trans-Golgi Network of CHO-K1 Cells. Biochemical Journal 1999, 342 (Pt 3), 633. 10.1042/0264-6021:3420633.

(33) Wang, Z.-H.; Xia, Y.; Wu, Z.; Kang, S. S.; Zhang, J.; Liu, P.; Liu, X.; Song, W.; Huin, V.; Dhaenens, C.-M.; Yu, S. P.; Wang, X.-C.; Ye, K. Neuronal ApoE4 Stimulates C/EBPβ Activation, Promoting Alzheimer’s Disease Pathology in a Mouse Model. Prog Neurobiol 2022, 209, 102212. 10.1016/j.pneurobio.2021.102212.

(34) Sun, Y.; Islam, S.; Gao, Y.; Nakamura, T.; Zou, K.; Michikawa, M. Apolipoprotein E4 Inhibits γ-Secretase Activity via Binding to the γ-Secretase Complex. J Neurochem 2022. 10.1111/jnc.15750.

