## Supplementary Figures 1 and 2 for "Apolipoprotein-based regulation of ganglioside metabolism upon secretase activity inhibition in iPSC-derived cerebral organoids"

**Figure S1. Ceramides and gangliosides levels upon treatment with secretase inhibitor in the ApoE3/3 AD cerebral organoids.** The initial precursors of gangliosides, namely dihydroceramide and ceramide, showed significant downregulation. However, this change was not reflected further in major ganglioside species. Each box plot indicates the minimum and maximum with a median of 10 values. ****P < 0.0001, ***P < 0.001, **P < 0.01, *P < 0.05 (Groups were compared using two-tailed t-test).

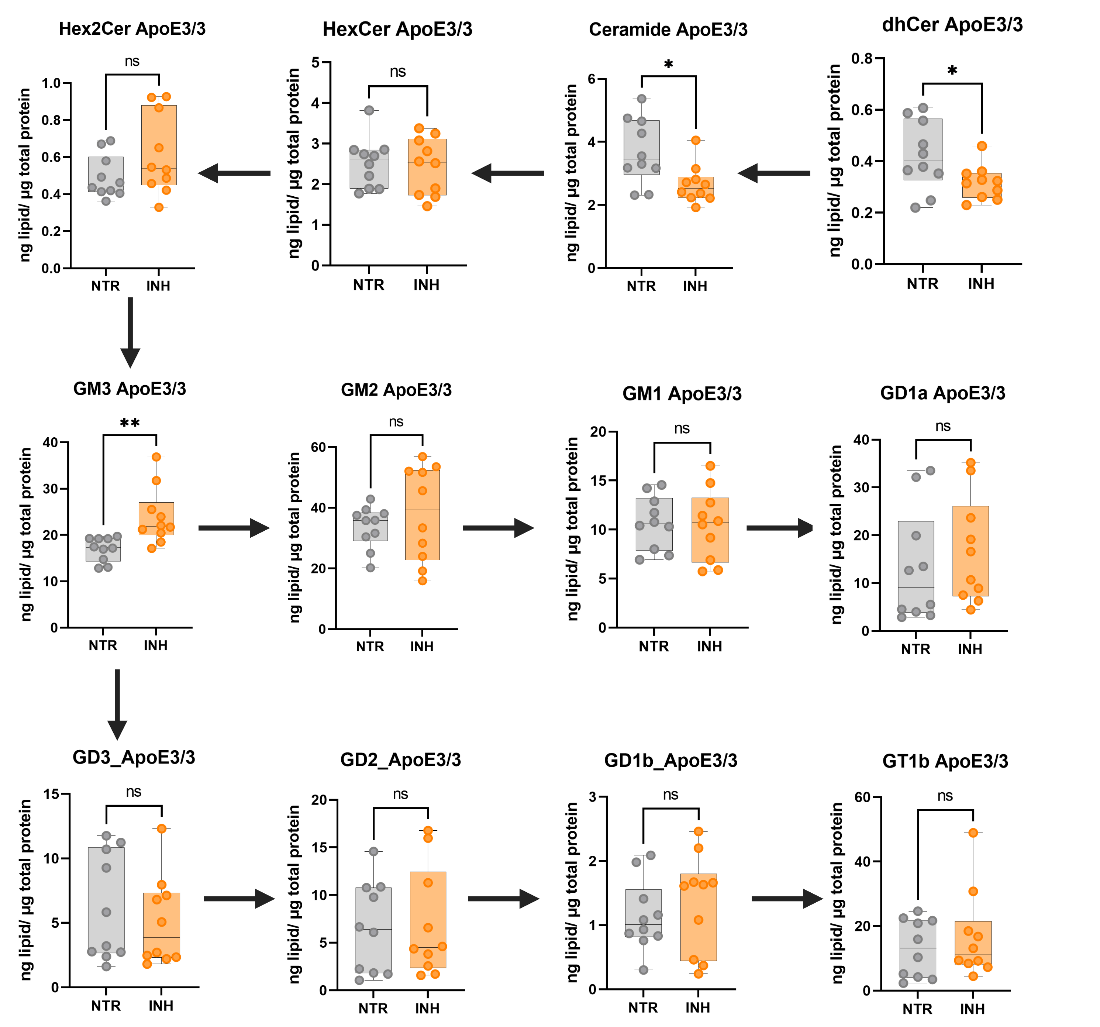

**Figure S2. Ceramides and gangliosides levels upon treatment with secretase inhibitor in the ApoE4/4 AD cerebral organoids.** We detected an upregulation in the biosynthetic pathway of gangliosides, including its precursors dihydroceramide, ceramide, glucosylceramide and lactosylceramide species. Each box plot indicates the minimum and maximum with a median of 10 values. ****P < 0.0001, ***P < 0.001, **P < 0.01, *P < 0.05 (Groups were compared using two-tailed t-test).

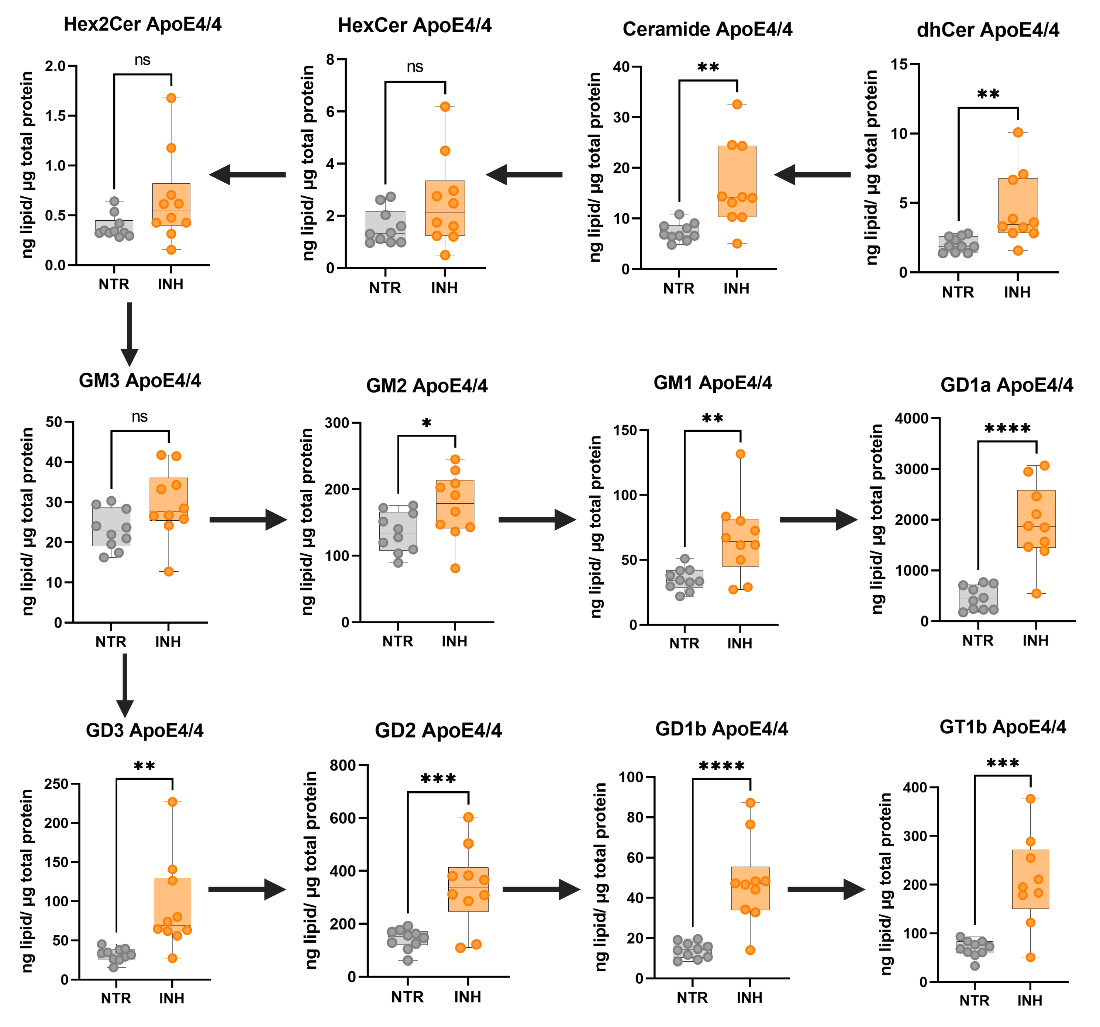
