## Supplemental Table 1 for "Apolipoprotein-based regulation of ganglioside metabolism upon secretase activity inhibition in iPSC-derived cerebral organoids"

**Table S1.** Selected reaction monitoring (SRM) library of precursor/product ion transitions for the analysis of gangliosides in positive and negative ion detection modes

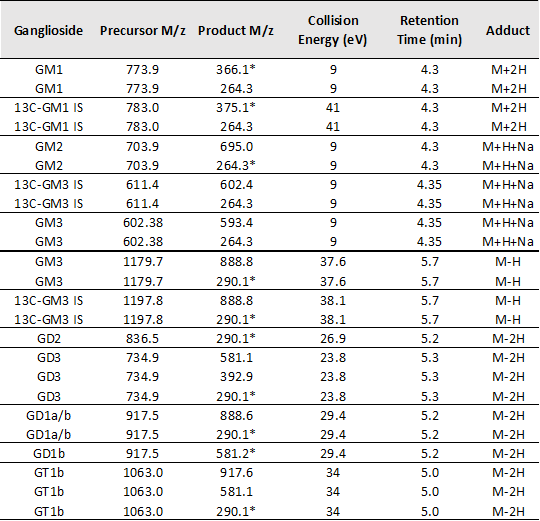

*Transitions used for the quantitation of the respective gangliosides.
